# Integrated single cell and bulk gene expression and ATAC-seq reveals heterogeneity and early changes in pathways associated with resistance to cetuximab in HNSCC sensitive cell lines

**DOI:** 10.1101/729384

**Authors:** Luciane T. Kagohara, Fernando Zamuner, Michael Considine, Jawara Allen, Srinivasan Yegnasubramanian, Daria A. Gaykalova, Elana J. Fertig

## Abstract

Identifying potential mechanisms of resistance while tumor cells still respond to therapy is critical to delay acquired resistance. We generated the first comprehensive multi-omics, bulk and single cell data in sensitive head and neck squamous cell carcinoma (HNSCC) cells to identify immediate responses to cetuximab. Two pathways potentially associated with resistance were focus of the study: regulation of receptor tyrosine kinases through the transcription factor TFAP2A, and epithelial-to-mesenchymal transition (EMT) process. Single cell RNA-seq demonstrates heterogeneity, with cell specific *TFAP2A* and *VIM* expression profiles in response to treatment. RNA-seq and ATAC-seq reveal global changes within five days of cetuximab therapy, suggesting early onset of mechanisms of resistance; and corroborates cell line heterogeneity, with different TFAP2A targets or EMT markers affected by therapy. Lack of *TFAP2A* reduces HNSCC growth and is enhanced by cetuximab and JQ1. Regarding the EMT process, short term cetuximab therapy has the strongest effect on inhibiting migration. *TFAP2A* silencing does not affect cell migration, supporting an independent role for both mechanisms in resistance. Overall, we show that immediate adaptive transcriptional and epigenetic changes induced by cetuximab are heterogeneous and cell type dependent; and independent mechanisms of resistance arise while tumor cells are still sensitive to therapy.

## INTRODUCTION

Cancer targeted therapies are designed to block specific relevant pathways for tumor progression. By doing so, these agents inhibit tumor growth resulting in prolonged survival ^1^. However, these therapies are not curative due to acquired resistance that develops within a few years of therapy ^2^. The mechanisms behind the tumor evolution from responsive to resistant state are not fully understood ^3,4^ but can involve mutations to the gene targeted, activation of downstream genes, and activation of alternative pathways ^5^. Studies aiming to characterize the mechanisms of resistance have shown an important role of tumor heterogeneity and from cell adaptive responses to these therapies as the sources of resistance ^6^. The presence of a multitude of cell clones increases the chances of the existence of intrinsic resistant tumor cells that are selected and will keep growing despite the treatment ^6^. In addition, sensitive cell clones have the ability of activating alternative pathways to overcome the blockade of the targeted growth pathway ^7^. Investigating the relevant early adaptive mechanisms that are potential drivers of resistance is critical to introduce early alternative therapies before the phenotype evolves as the dominant feature among the cancer cells.

Currently, cetuximab is the only FDA approved targeted therapeutic for HNSCC ^8^ and was selected based on pervasive overexpression of EGFR and its associations with outcomes in HNSCC ^9,10^. As with other targeted therapies, virtually all HNSCC patients develop acquired resistance limiting its clinical application ^11^. The near universal emergence of resistance and intermediate time rate at which it occurs mark cetuximab treatment in HNSCC as an ideal model system to study resistance. Little is known about the immediate transcriptional and epigenetic changes induced by cetuximab in the very early stages of therapy. We and others have found that compensatory growth factor receptor signaling regulated by *TFAP2A* and EMT, both associated with resistance, are altered while cells are still sensitive to therapy ^12,13^. Therefore, their precise role in resistance and timing at which they induce phenotypic changes remains unknown. It is critical to isolate the timing and effect of each of these pathways during cetuximab response to delineate their subsequent role in resistance.

We hypothesize that the up-regulation of mechanisms of resistance arise while HNSCC cells are still sensitive to cetuximab and that some of these mechanisms are associated with chromatin remodeling induced as an immediate response to therapy. Our previous study showed in vitro up-regulation of *TFAP2A* one day after treatment with cetuximab ^12^. Together with the fact that some of its targets are receptor tyrosine kinases ^14,15^, it is very probable that this is one of the mechanisms activated by HNSCC cells to overcome EGFR blockade and that will induce resistance. Schmitz et al. ^13^ also demonstrated that mechanisms of resistance to cetuximab arise early in the course of HNSCC patients therapy by detecting EMT up-regulation after only two weeks treatment. The stimulation of the EMT phenotype is a common mechanism of resistance to different cancer therapies, including cetuximab ^16–18^. In the current study, we focused on these two pathways to investigate how the transcriptional and epigenetic status are rewired while cancer cells are still sensitive to cetuximab.

In order to verify our hypothesis, we performed single cell RNA-sequencing (scRNA-seq) to understand how three HNSCC cell lines and each of their clones respond to a short time course cetuximab therapy. Then, using bulk RNA-sequencing (RNA-seq) and ATAC-seq we investigated the changes of two relevant pathways (TFAP2A and EMT). We verified the heterogeneous response to cetuximab among the cell models with cell line specific adaptive responses to cetuximab and clear disturbances in both pathways. *TFAP2A* regulates HNSCC growth in vitro and in its absence cells proliferate less. A potential interplay with the EMT was not verified, suggesting that two independent resistance mechanisms to cetuximab are early events in the course of therapy. The response to the combination therapy cetuximab and JQ1, although heterogeneous, is more efficient to cell growth control than anti-EGFR therapy alone suggesting that combined therapies blocking multiple growth factors are beneficial in the early stages of therapy.

## RESULTS

### TFAP2A and EMT expression are heterogeneous among cell lines

To investigate the heterogeneous responses induced by therapy before resistance developed, sensitive HNSCC models were used to interrogate the immediate changes induced by cetuximab. Based on previous work demonstrating HNSCC cell lines sensitivity to cetuximab ^16,30^ and confirmed by proliferation assay, we chose the cell lines SCC1, SCC6 and SCC25 (**Supplementary Figure 1**). To verify heterogeneity and how each of the cell clones respond to cetuximab, we performed single cell RNA-seq (scRNA-seq). The cell lines received cetuximab (treated) or PBS (untreated) and after a total of five days the cells were collected in single cell suspensions for the library preparations and sequencing (**Figure 1A**).

**Figure 1.**
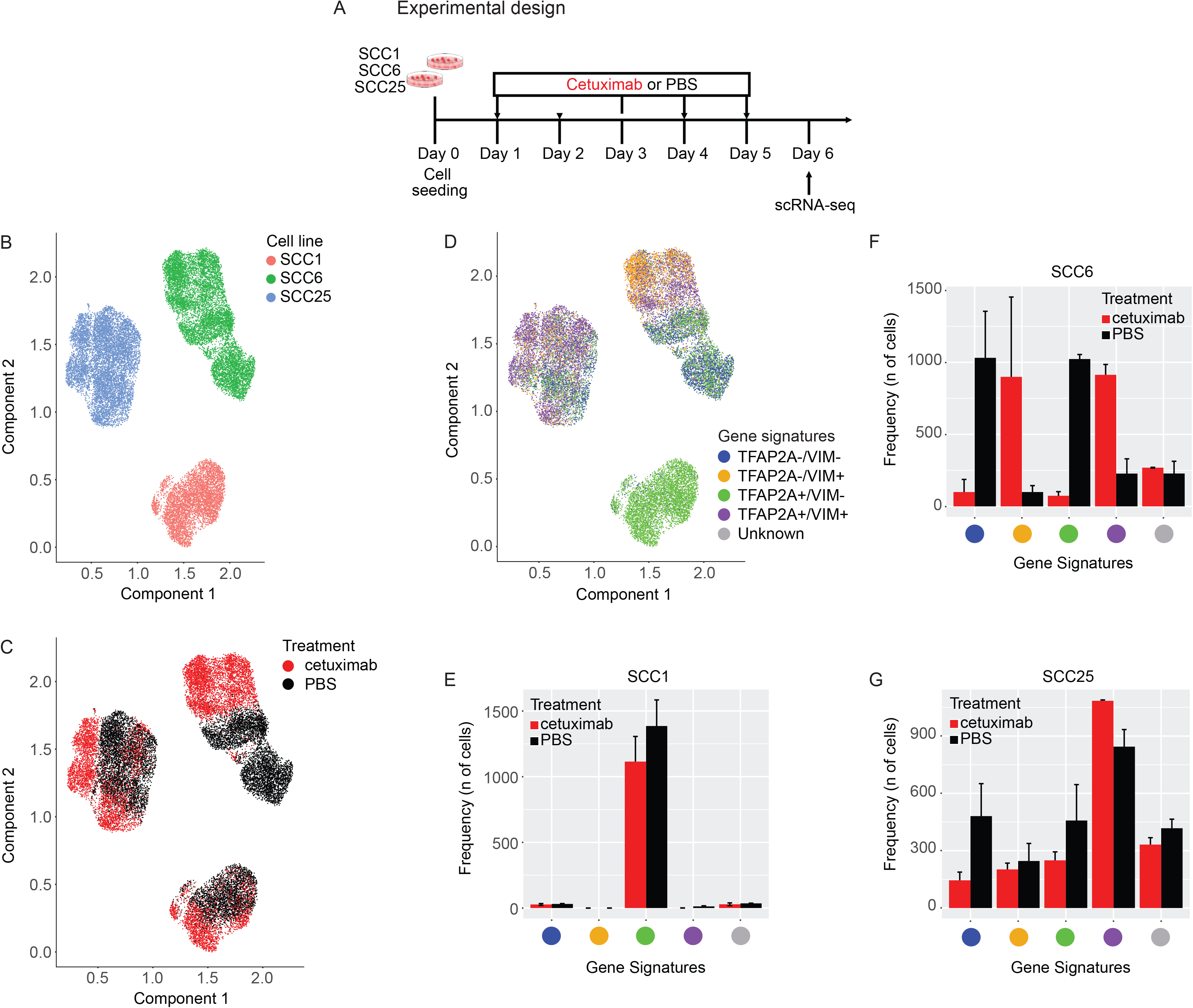
**(A)** SCC1, SCC6 and SCC25 cell lines were treated with cetuximab or PBS (untreated controls) for five consecutive days after which cells were collected for single cell RNA-seq (scRNA-seq). **(B)** scRNA-seq analysis demonstrates that each cell line present a specific gene expression profile. **(C)** In response to cetuximab, the SCC6 treated (red) and untread (black) clones separate completely while the SCC1 and SCC25 present some overlap in the distribution regarding the transcriptional profile. **(D)** Inter-cell heterogeneity is more evident for *TFAP2A* and *VIM* mRNA levels, with SCC1 presenting high levels of *TFAP2A* and no expression of *VIM*. The co-expression analysis shows that in SCC1 there is no change in the levels of *TFAP2A* or *VIM* in response to cetuximab; SCC6 treated cells are *VIM+* (orange and purple) while untreated are negative (green and blue) with different status for *TFAP2A* expression; and most of the SCC25 cells responding with increase in *VIM* but with some untreated clones presenting the same expression profile for *VIM* and *TFAP2A* (purple) and with *VIM-* clones only detected in the untreated group. **(E, F, G)** Bar plots represent the number of treated and untreated cells per each gene signature.

Based on the whole transcriptomic profile, each cell line cluster completely separate from each other (**Figure 1B**) demonstrating expected inter-cell line heterogeneity. Analyzing the cell clusters according to cetuximab therapy, we noted that each cell line presents specific early transcriptional responses. There is a clear separation between treated and untreated cells in SCC6 (**Figure 1C**), suggesting that in only five days anti-EGFR therapy induces significant transcriptional changes when compared to the untreated (PBS) cells. For SCC1 and SCC25, there are treated cells that cluster with the untreated ones (**Figure 1C**) and most probably in these cell lines prolonged exposure is necessary for more significant changes in gene expression.

To investigate the immediate emergence of potential mechanisms of resistance, we investigated the expression of *TFAP2A* and *VIM*, alone or concomitantly, to verify the behavior of these pathways (transcription regulation by TFAP2A and EMT process) in response to cetuximab. We evaluated the expression of *TFAP2A* and *VIM* genes in the individual cells (**Supplementary Figure 2**) and used the individual markers expression levels to classify each individual as double negative (*TFAP2A-/VIM-*), *TFAP2A* positive (*TFAP2A+/VIM-*), *VIM* positive (*TFAP2A-/VIM+*) and double positive (*TFAP2A+/VIM+*) (**Figure 1F**). The scRNA-seq analysis of the three cell lines show heterogeneity regarding the expression of *TFAP2A* and *VIM* genes. Cetuximab treated and untreated SCC1 show high levels of *TFAP2A* and absence of *VIM* expression (**Supplementary Figure 2**, **Figure 1D and E**) suggesting no influence of therapy in these two markers for this specific cell line. SCC6 cells present a definite shift in the expression of *VIM* with the anti-EGFR blockade, with untreated cells presenting down-regulation when compared to the treated cells. The shift in *VIM* expression was independent of the *TFAP2A* status (**Supplementary Figure 2**, **Figure 1D and F**), without apparent variation in the proportions of positive and negative cells in response to cetuximab. Interestingly, the majority of SCC25 cells are double positive with or without cetuximab therapy. In the presence of EGFR blockade, *VIM* expression is positive among most of the treated clones (double positive), while the proportions of untreated SCC25 cells expressing or lacking *VIM* are approximate (**Supplementary Figure 2, Figure 1D and G**).

Based on the SCC6 expression profile, there is evidence that cetuximab is capable of inducing *VIM* expression and, corroborating the observation from Schmitz et al. (11), that cetuximab induces EMT markers early on in the course of therapy. However, most of the transcriptional changes in response to cetuximab are cell type dependent.

### Cetuximab induces immediate gene expression changes in HNSCC in vitro

In order to evaluate the timing of the changes in the TFAP2A targets and EMT markers and to interrogate each of the pathway genes individually, we performed daily measurements in treated and untreated groups for all three cell lines with bulk RNA sequencing (RNA-seq) (**Figure 2A**).

**Figure 2.**
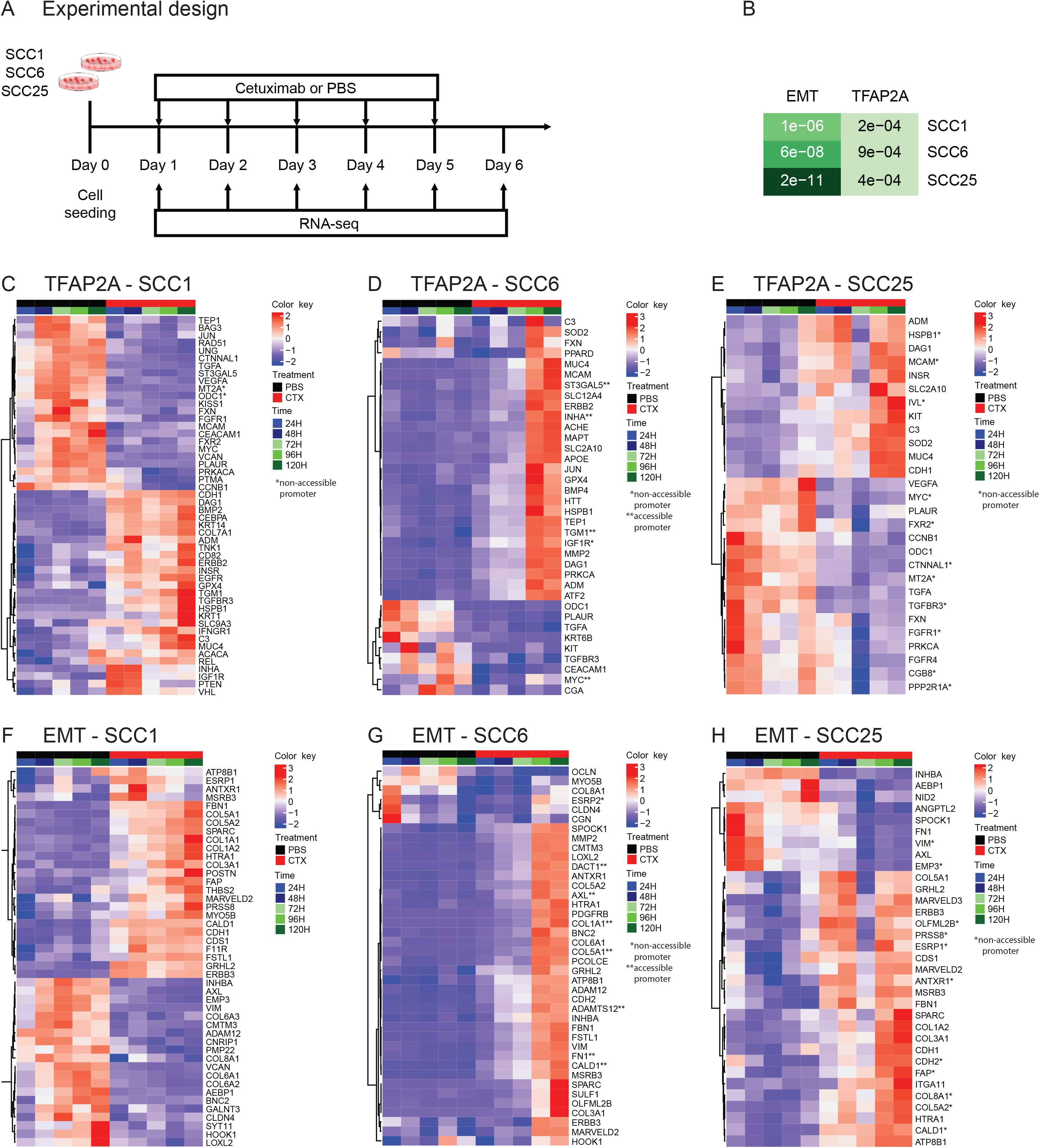
**(A)** SCC1, SCC6 and SCC25 cell lines were treated with cetuximab or PBS (untreated controls) for five consecutive days and cells were collected daily for bulk RNA-seq (RNA-seq). **(B)** Among the genes differentially expressed among all three cell lines as a response to cetuximab therapy the gene set enrichment analysis show significant presence (p≤0.05) of genes that are TFAP2A targets or that participate in the EMT process. When analyzed individually, the TFAP2A and EMT differential expressed genes are specific in each of the cell lines. **(C and F)** SCC1 and **(E and H)** SCC25 present changes as soon as 24 hours (1 day) after cetuximab therapy, while in **(D and G)** SCC6 the changes are only detected at 96 hours (4 days) after cells are treated.

Transcriptional changes induced by cetuximab can be detected genome wide almost immediately after therapy. Differential expression analysis of all time points indicate that hundreds of genes have their transcriptional profile changed as a response to anti-EGFR therapy in all three HNSCC cell lines with changes occurring as early as 24 hours after treatment (**Supplementary Figure 3A**). In order to investigate the changes in the activity of TFAP2A transcription factor, we followed the expression of its targets identified using the TRANSFAC database (12,13). To analyze the status of the EMT pathway, we analyzed the EMT markers from the gene signature described by Byers et al. that can predict resistance to anti-EGFR and anti-PI3K therapies (27). When each cell line was investigated separately, the gene set enrichment analysis comparing cetuximab and untreated timepoints showed that among the differentially expressed genes in SCC1, 55 are TFAP2A targets (p=2.2e-04) and 49 are EMT markers (p=1.1e-04); in SCC6, there are 46 genes from each pathway (TFAP2A p=9e-04, EMT p=6e-08); and in SCC25, there are 40 TFAP2A targets (p=4.3e-04) and 46 EMT markers (p=2.2e-11) (**Figure 2B**). Although there was no variation in the expression of *TFAP2A* and *VIM* in SCC1, there are still significant changes to other markers in both pathways that are potentially associated with future development of acquired cetuximab resistance. The cell lines SCC1 and SCC25 present immediate transcriptional changes to the cetuximab therapy and most of the genes present expression changes in the first 24 hours of therapy (**Figure 2C, F, E and H**). SCC6 transcriptional response to anti-EGFR treatment takes longer and most of the changes are noticeable after 96 hours of therapy (**Figure 2D and G**), which is in agreement with the observed behavior of this cell line to the cetuximab therapy (**Supplementary Figure 1**).

### Chromatin changes can be detected early in the course of cetuximab therapy in vitro

We hypothesized that epigenetic rewiring induced by cetuximab is the most probable cause of the adaptive transcriptional changes we detected with RNA-seq. To verify if chromatin remodeling occurs early during cetuximab treatment and if it affects the TFAP2A targets and EMT genes, we measured global chromatin accessibility by ATAC-seq in cells treated with cetuximab and in the untreated controls after five days of therapy (**Figure 3A**).

**Figure 3.**
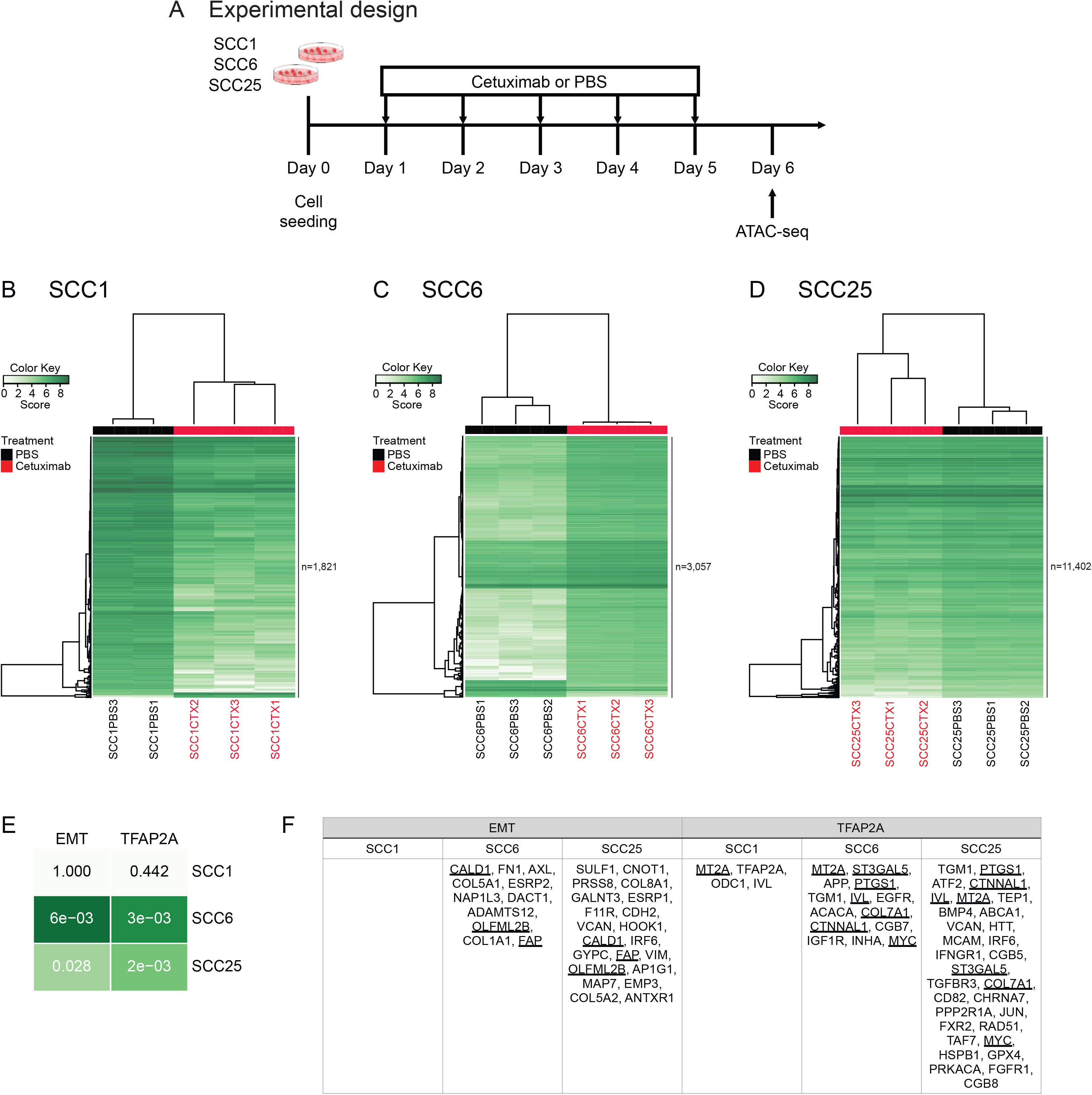
**(A)** ATAC-seq was performed after SCC1, SCC6 and SCC25 were treated for 5 days with cetuximab and also in the untreated (PBS) controls. **(B, C and D)** Differential binding analysis show that the promoters accessibility changes in response to 5 days of therapy are capable of separating the cetuximab from the PBS replicates in all three cell lines. **(E)** With the exception of SCC1, there are enrichment for TFAP2A and EMT promoters among the ATAC-seq peaks in SCC6 and SCC25. **(F)** The differential binding analysis show that SCC25 is the gene with the highest number of genes with chromatin changes in response to cetuximab and also identified promoters that are changed in more than one cell line (underlined gene names).

Cetuximab induces significant chromatin changes after only five days of therapy (**Supplementary Figure 3B**). Differential bound analysis, to identify the accessible protein-DNA binding regions in cetuximab versus untreated groups, show that there are a total of 1,690 binding regions, common to SCC1, SCC6 and SCC25, that have their structure changed as a response to therapy. The unsupervised clustering of these common regions separate the samples that were treated from the untreated controls (**Supplementary Figure 3B**). These findings suggest that epigenetic rewiring is an early event in response to cetuximab and is probably involved in the regulation of some relevant transcriptional changes observed.

The differential binding analysis was performed for each cell line individually to identify cell specific chromatin changes in response to cetuximab (**Figures 3B, C and D**). Each of the three cell lines presents specific chromatin changes that separate the groups of treated and untreated replicates. SCC1 and SCC6 show significant promoters reconfiguration as a response to therapy with 1,821 and 3,057 sites remodeled, respectively (**Figures 3B and C**). SCC25 presents the largest number of gene promoters remodeled with 11,402 promoter binding sites (including genes with more than one binding site) as a result of short term therapy (**Figure 3D**).

The gene set enrichment analysis identified genes from the TFAP2A and EMT pathways in the list of promoters that have chromatin structural changes induced by cetuximab. Promoter region reconfiguration during cetuximab treatment in the SCC1 cell line was detected in only four genes from the TFAP2A pathway and no changes in EMT promoters is present (**Figure 3E and F**). Suggesting that in this cell line the transcriptional changes in both pathways are not regulated by chromatin remodeling. A total of eleven promoters from the TFAP2A pathway (p=3e-03, **Figure 3E and F**) and the same number of EMT gene promoters (p=6e-03, **Figure 3E and F**) have their chromatin structure changed by the anti-EGFR therapy in SCC6. The SCC25 cell line presents, as a response to cetuximab, chromatin changes in 31 TFAP2A pathway (p=0.028, **Figure 3E and F**) and in 21 EMT promoters (p=2e-03, **Figure 3E and F**). Interestingly, all chromatin changes to the SCC25 binding sites make them less accessible when compared to the untreated controls. The ATAC-seq findings suggest that even after a short time exposure of HNSCC cells to cetuximab in vitro, genes from pathways that are associated with acquired resistance present remodeling that could potentially result in altered transcription factors binding.

The genes with transcriptional and chromatin alterations in response to short time treatment with cetuximab are marked with one (non-accessible after cetuximab) or two stars (accessible after cetuximab) in the RNA-seq heatmaps in Figure 2. As would be expected, the correlation between accessibility and expression is not true for all genes. Although a few relevant genes, such as *AXL* (**Figure 2D**), known to be up-regulated in acquired resistance to different targeted agents, presents open chromatin combined with up-regulation in SCC6 treated cells.

### TFAP2A controls HNSCC proliferation in vitro

The role of *TFAP2A* in HNSCC is poorly characterized. As a transcription factor, it is capable of regulating the expression of several growth factor receptors (EGFR, HER2, TGFBR3, FGFR1, IGFR1 and VEGF) 12. In order to investigate the role of *TFAP2A* in HNSCC cell proliferation in vitro, we used siRNA assay for gene silencing and measured growth rates for five days following therapy (**Figure 4A**). All transfected cell lines present lower growth rates when compared to the parental cell lines (**Figure 4B, C and D**, black full and dashed lines). The effect of *TFAP2A* is more prominent in SCC1 and SCC25 if compared to SCC6. This is probably related to the fact that both cell lines present *TFAP2A* expression in most of the cell clones, as shown by the scRNA-seq (**Figure 1D**).

**Figure 4.**
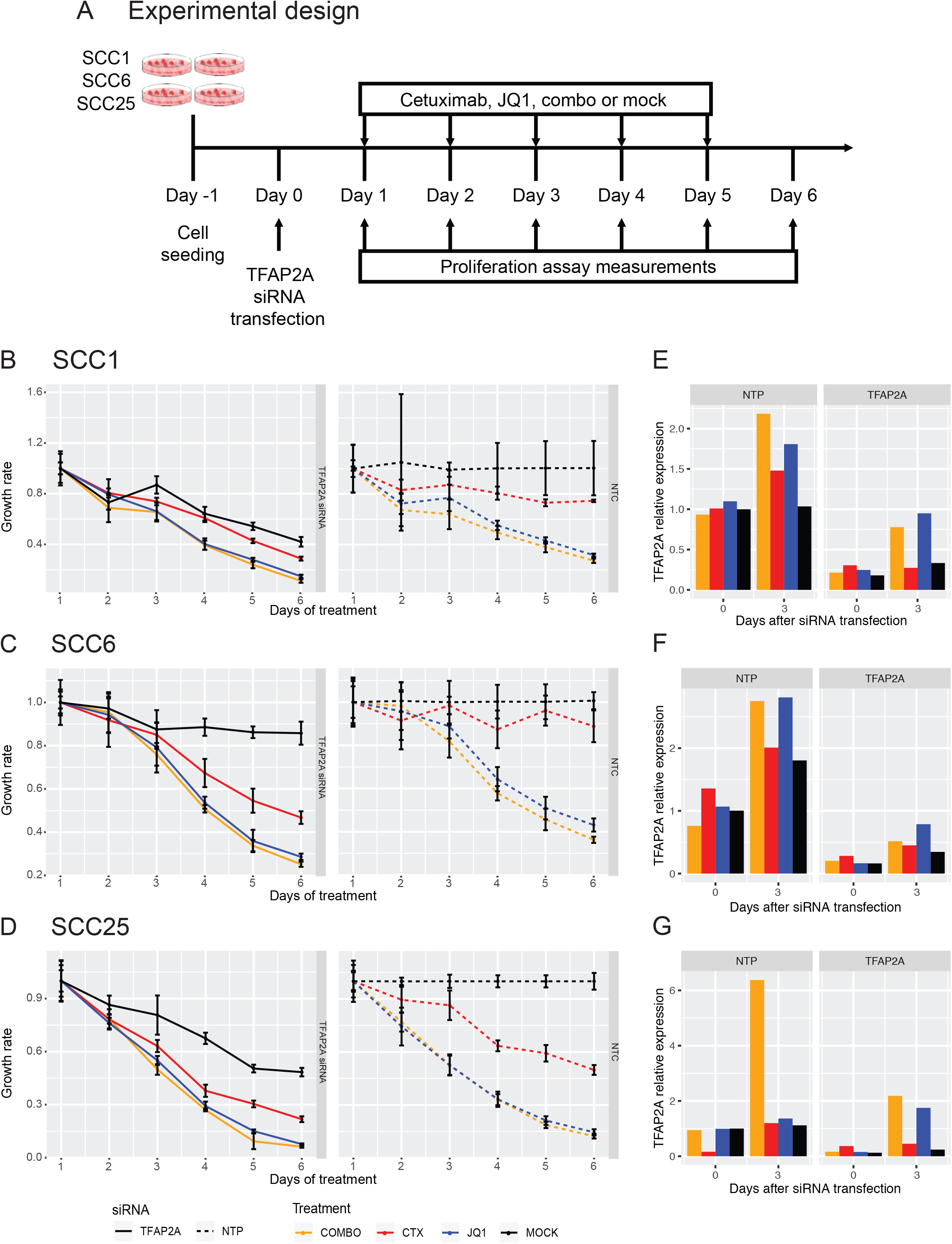
**(A)** Functional validation of the role of *TFAP2A* in HNSCC in vitro was evaluated by siRNA gene silencing in SCC1, SCC6 and SCC25. Cells were treated with cetuximab, JQ1, combination (combo) or vehicle (mock) for five days and the impact of gene knockdown and therapy was determined by measuring proliferation rates. **(B, C and D)** Transfected groups (full lines, left) were compared to the groups with normal levels of TFAP2A (dashed lines, right - NTC). In all cell lines, TFAP2A knockdown induce lower proliferation rates (black lines) at different levels depending on the cell. Cetuximab treatment (red lines) present a synergistic effect but JQ1 (blue lines) efficacy is even greater in reducing cell growth. Little effect is noted with the combination (orange lines, COMBO) when compared to the effects of JQ1 alone.

Combined with the effects of *TFAP2A* transient knock-down, we investigated the role of cetuximab and JQ1 on HNSCC growth. JQ1 is a bromodomain inhibitor, that blocks the transcription of cell growth regulators (e.g., c-Myc) and multiple RTKs, and was previously shown to delay acquired cetuximab resistance 31. Cetuximab or JQ1 was added to cell culture media once cells were transfected with *TFAP2A* siRNA, and proliferation was measured daily (**Figure 4A**). We also verified how cells would respond to the combination (combo) of both drugs in vitro (**Figure 4A**).

Cetuximab therapy potentiates growth inhibition in the absence of *TFAP2A* (**Figure 4 B, C and D**, red full and dashed lines) with synergistic effect potency dependent on the cell line. SCC1 presents very similar *TFAP2A* expression in treated and untreated cell clones (**Figure 1D**), and the effect of gene knockdown with anti-EGFR therapy is not as significant as observed in SCC6 and SCC25. A stronger effect on proliferation control was observed with JQ1 treatment (**Figure 4 B, C and D**, blue full and dashed lines), most probably due to the silencing of another proliferation factor (c-Myc) and/or RTKs. Interestingly, the combination therapy of cetuximab and JQ1 does not provide a significantly stronger synergistic effect (**Figure 4B, C and D**, orange full and dashed lines). *TFAP2A* transient knockdown was confirmed by qRT-PCR (**Figure 4E, F** and **G**). These results indicate that in HNSCC in vitro, the transcription factor TFAP2A is an essential regulator of cell growth.

### Cetuximab inhibits HNSCC cell migration in vitro

To investigate the role of cetuximab and JQ1 in the EMT pathway, we performed the scratch assay on SCC1, SCC6 and SCC25 cells treated with both drugs alone or in combination. The cells were seeded in cell migration inserts (Ibidi) and treatment with cetuximab, JQ1, combo or vehicle (mock) 48 hours later. Once confluence was reached (72 hours after seeding) the insert was removed, and gap closure was measured at 0, 6, 12 and 24 hours (**Figure 5A**).

**Figure 5.**
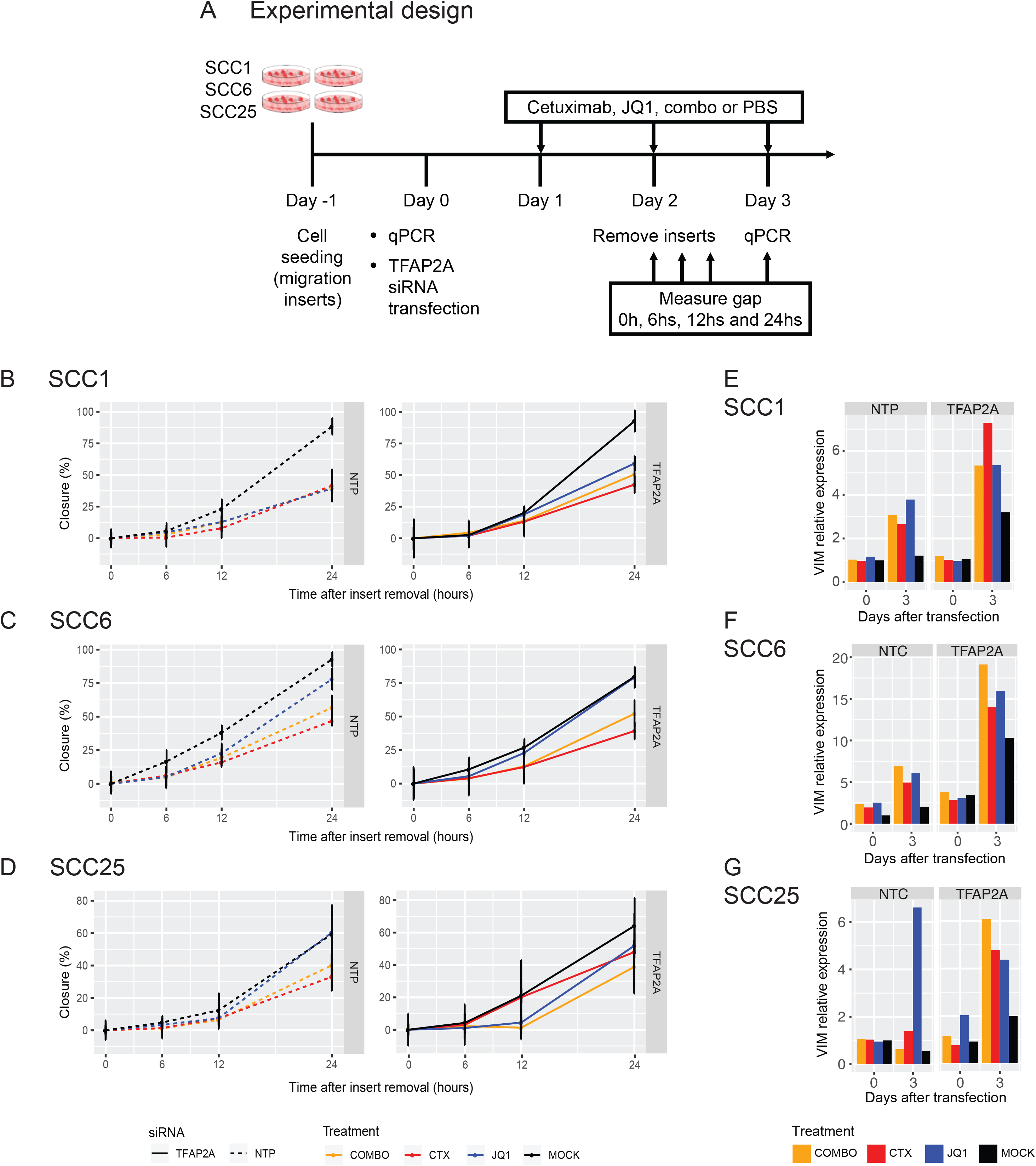
**(A)** To further evaluate the interplay between *TFAP2A* and EMT, cells transfected with siRNA against *TFAP2A* and treated with cetuximab, JQ1, combination (combo) or vehicle (mock) were used for a migration assay. Migration was measured for a total of 24 hours immediately after insert removal. **(B, C and D)** No significant changes in migration was noted when comparing the non-transfected (dashed lines, left) and transfected (full lines, right - NTP) SCC1, SCC6 and SCC25 cells and different treatment groups. **(E, F and G)** Although migration changes were not observed, there are changes in VIM expression as response to siRNA silencing and the different therapies in all three cell lines.

Cetuximab treatment resulted in cell migration inhibition in all three cell lines (**Figure 5B, C** and **D**) when compared to the corresponding untreated cells. The treatment with JQ1 had distinct effects in each of the cell models. Migration of SCC1 with cetuximab, JQ1 or combined therapy did not present any change and the inhibition effects were the same for all treatment groups when compared to the untreated cells (**Figure 5B**). In SCC6, therapy also suppressed migration relative to the absence of treatment. SCC6 cells treated with JQ1 migrate faster than in the presence of cetuximab while the combination therapy reduces migration but not as efficiently as cetuximab monotherapy (**Figure 5C**). Although JQ1 was able to reduce SCC1 and SCC6 migration, there was no effect on the migratory abilities of SCC25 and the cells maintain the same rate as untreated cells. Cetuximab had the strongest effect on repressing SCC25 migration and the combination also reduced motility in a lower extent (**Figure 5D**).

There is no reference in the literature to a possible interplay between the *TFAP2A* and EMT genes in HNSCC. Since transcriptional factors regulate multiple targets, we also investigated this potential interaction. HNSCC cell lines migration is not impacted by the lack of *TFAP2A* after transfection with siRNA. SCC1, SCC6 and SCC25 transfected with *TFAP2A* siRNA (**Figure 5E, F** and **G**) present the same migration rates as the non-transfected cell lines (**Figure 5B**). The different therapies inhibit migration in a cell type specific manner (**Figure 5E, F** and **G**). The scratch assay observations suggest that *TFAP2A* do not directly regulate EMT genes as silencing of the transcription factor does not affect migration directly. The effects in migration are only associated with cetuximab or JQ1 therapy.

## DISCUSSION

Approximately 90% of HNSCC present high expression of EGFR protein, and cetuximab seemed to be a reasonable targeted therapy for these tumors (28). However, just a small fraction of patients respond to cetuximab, and virtually all responders develop acquired resistance (29). To prolong disease control, it is crucial to identify the changes related to resistance while the tumor is still responsive to cetuximab. Currently, there are no biomarkers to predict the drug response, and the mechanisms of resistance are poorly characterized in HNSCC (30,31). In a recent time course study to investigate the transcriptional and DNA methylation signatures driving acquired cetuximab resistance in HNSCC, we found that an essential driver of resistance to anti-EGFR targeted therapies, FGFR1, is epigenetically regulated during chronic exposure to cetuximab and provide strong evidence that epigenetic alterations can drive acquired resistance (10).

Using a single cell and bulk multi-omic approach, we investigated the early responses to cetuximab in HNSCC in vitro models to identify the gene expression and epigenetic mechanisms that are potential drivers of resistance. Treating three HNSCC cell lines for a short period of time, we were able to demonstrate that transcriptional and chromatin rewiring are early events as a response to therapy and that they happen globally and include genes previously described to be involved in resistance to cetuximab. Here, we investigated three HNSCC cell lines (SCC1, SCC6 and SCC25) and their responses to cetuximab in the first days of therapy. scRNA-seq analysis demonstrates that even the untreated cells demonstrate specific transcription profiles that prove inter-cell lines heterogeneity. We observed that meanwhile *VIM* expression presents a shift after cetuximab therapy in SCC6 and SCC25, it does not have the same behavior in SCC1.

We further performed a short time course experiment to measure daily the transcriptional changes induced by cetuximab in the three cell lines to verify the cell specific changes to the TFAP2A targets and EMT genes. Although we did not observe changes in *TFAP2A* and *VIM* in SCC1 at the single cell level, other genes from these pathways are altered as soon as 24 hours after treatment initiation suggesting that other markers respond with changes in expression to cetuximab. Each cell line presents specific changes to distinct genes from the pathways interrogated. SCC1 and SCC25 present changes after only 24 hours of therapy while in SCC6 those changes are noticed within 96 hours of therapy. These results reflect the initial observation in growth rates under cetuximab therapy, where SCC6 presents a resistant-like behavior with decreased proliferation only after 96 hours under cetuximab (**Supplementary Figure 1**) or stable slower growth with therapy (**Figure 4C**). We have previously observed that anti-EGFR targeted therapy in vitro is capable of inducing immediate transcriptional changes in the HaCaT keratinocyte cell line model with constitutive *EGFR* activation (10). Here we corroborate this observation by showing that two HNSCC cell lines also present immediate changes to cetuximab and in pathways relevant for resistance. Altogether, these are evidence that adaptive responses to targeted therapies can occur to genes that are involved in driver pathways of resistance and while cancer cells still sensitive to the therapy.

In another study, we have shown that while SCC25 acquire cetuximab resistance due to chronic exposure (32), the transcriptional changes occur a few weeks prior to the promoter hyper- or hypomethylation, with the latter been detected when cells are already resistant. Here, we investigated the hypothesis that some of the genes involved in resistance are controlled by chromatin remodeling that occurs prior to methylation, while the cells are still sensitive to the therapy, and drive a proportion of the expression changes. After five days of anti-EGFR blockade, chromatin structure differs between cetuximab and untreated groups in the three HNSCC cell lines as shown by ATAC-seq. We hypothesize that the events that result in acquired resistance go from chromatin changes in the early stages of cetuximab therapy and reflect in transcriptional changes to overcome EGFR inhibition, and that are finally stabilized by gain or loss of methylation. It was previously shown in vitro that *CDKN2A* silencing initially happens through histone modifications leading to loss of gene expression, followed by promoter methylation to lock the repressive state (33). Our findings, together with Bachman et al., suggest that while chromatin rewiring results in gene expression changes, this epigenetic state is still reversible and require DNA methylation to be maintained and inherited. It is critical to determine the timing in treatment that reversible epigenetic alterations develop to allow alternative therapies to be effective. Short term exposure to targeted therapies can induce reversible chromatin changes that will lead to resistance, while chronic exposure induces DNA methylation changes that are more steady and observed in stable resistant states (34).

*TFAP2A* encodes a transcription factor that binds to growth factor receptors and is most probably up-regulated to overcome the lack of EGFR activity. One proof that this is a potential mechanism of resistance is our previous observation that as a response to anti-EGFR therapy, *TFAP2A* mRNA level is up-regulated with only 24 hours of therapy initiation in vitro (10). The TFAP2A transcription factor has dual-function and can play a role as a tumor suppressor gene (transcriptional repressor) or oncogene (transcriptional activator) depending on the tumor type. Although a previous study showed that in vitro down-regulation of *TFAP2A* in HNSCC is associated with decreased proliferation (35), another study pointed to the same direction as our findings. In nasopharyngeal carcinoma, *TFAP2A* silencing in vitro and in vivo results in slower cancer cell proliferation and that patients with high tumor levels of the gene present poorer survival compared to those with lower expression (36). *TFAP2A* up-regulation is a feature of other tumor types such as neuroblastoma, pancreatic cancer and acute myeloid leukemia ^32–34^. In our in vitro models, *TFAP2A* knock-down resulted in slower cell growth showing the relevance of this transcription factor to HNSCC proliferation in vitro. This finding together with the observation that cetuximab has a synergistic effect is evidence that TFAP2A downstream targets could be new therapeutic markers for combination approaches that will result in prolonged disease control.

The EMT process has also been previously associated with acquired resistance to anti-EGFR targeted therapies in cells with mesenchymal phenotype ^7,18,35,36^. We found a significant number of EMT gene promoters among those undergoing remodeling after five days of therapy in SCC6 and SCC25. Among the EMT genes up-regulated by EGFR blockade are a few collagenases, most probably related to providing tumor cells ability to invade the extracellular matrix. One interesting finding is that the gene *AXL* is up-regulated after 96 hours of cetuximab therapy in the SCC6 cells and this is also correlated with a more accessible promoter. *AXL* is a receptor tyrosine kinase known to mediate resistance to cetuximab and is possibly an alternative mechanism HNSCC cells in vitro are activating to keep proliferating under therapy ^37–39^. This observation suggests that early chromatin modifications are involved in the development of acquired cetuximab resistance and that they can be detected in the beginning of the treatment.

Since the up-regulation of other RTKs, such as *AXL*, is a common finding in acquired anti-EGFR resistance, we tested a combination treatment with cetuximab to evaluate a possible synergistic effect on controlling cell growth more effectively than EGFR targeted therapy alone. JQ1 is a bromodomain inhibitor that preferentially binds to BRD4, a protein with high affinity for acetylated histone tails, which represses transcription of its targets ^40,41^. Among these target genes are RTKs known to be up-regulated as a resistance mechanism to anti-EGFR therapies ^31,42^. In this scenario, BRD4 inhibition seems a reasonable approach by acting as a “multi-targeted” therapy. Also, successful results in delaying acquired cetuximab resistance were shown when JQ1 or BRD4 knockdown were used in combination with cetuximab in HNSCC cell models or patient derived xenografts ^31^. In our short time course therapeutic model, we could not determine the time of resistance of development but we observed the inhibitory effects of JQ1 alone or in combination with EGFR blockade. JQ1 has stronger effect than cetuximab in controlling HNSCC proliferation in vitro and the addition of cetuximab has diverse impact in reducing cell growth depending on the cell line, with the strongest synergism observed in SCC6 cells. Although including cetuximab to the JQ1 therapy seems to have little effect on reducing proliferation, the combination probably has major impact on disease control by targeting various RTKs at the same time and delays acquired resistance due to reduction of alternative growth pathways tumor cells can use to overcome targeted inhibition. JQ1 is known to have a short half life reflecting in the necessity of elevated doses that would not be tolerated by cancer patients ^43,44^. Since there are currently other bromodomain inhibitors being evaluated in clinical trials with less toxicity than JQ1, further studies are necessary to identify one which would have a similar effect when combined with cetuximab in HNSCC.

Overall, our study demonstrates that transcriptional and chromatin changes induced by cetuximab therapy are early events that can be detected before acquired resistance develops. Here, we focused on two pathways, TFAP2A and EMT, previously described to be involved in resistance to cetuximab and other anti-EGFR therapies (10,11) Another major finding is how inter-cell heterogeneity can induce different changes to the same mechanisms of resistance to targeted therapies. Although we observe alterations in both TFAP2A and EMT pathways, the genes affected are different, and in one of the cell lines (SCC1), there is no apparent role of chromatin remodeling in the EMT transcriptional alterations. We demonstrate that two independent mechanisms of resistance present an early onset during the course of cetuximab therapy, suggesting that other mechanisms of resistance could also be deregulated. This observation is relevant since it demonstrates that to overcome resistance acquisition more than one combination therapy would be necessary. Alternatives like JQ1, that targets multiple drivers of resistance, are then promising and would allow the development of clinical trials or clinical decisions to be made without submitting patients to the expensive costs of genetic and genomic tests.

## MATERIAL AND METHODS

### Cell culture and proliferation assay

UM-SCC-1 (SCC1), UM-SCC-6 (SCC6) and SCC25 cells were cultured in Dulbecco’s Modified Eagle’s Medium and Ham’s F12 supplemented with 10% fetal bovine serum and maintained at 37°C and 5% CO_2_. A total of 25,000 cells were plated in quintuplicate in 6-well plates. Cetuximab (Lilly) was purchased from Johns Hopkins Pharmacy and JQ1 from Selleck Chemicals. Cell lines were treated daily with cetuximab (100nM), JQ1 (500nM), the combination, or vehicle (PBS+DMSO; mock) for five days. Proliferation was measured using alamarBlue assay (Thermo Scientific). AlamarBlue (10% total volume) was added to each well and fluorescence (excitation 544nm, emission 590nm) was measured after 4 hours of incubation at 37°C. A media only well was used as blank. The measurements were repeated in 3 independent experiments. Growth rate was calculated using the formula:

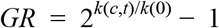

Where *k(0)* = fluorescence measured for non-treated cells and *k(c,t)* = fluorescence for treated cells^19^.

Parental cell lines were authenticated before and after all assays using short tandem repeat (STR) analysis kit PowerPlex16HS (Promega) through the Johns Hopkins University Genetic Resources Core Facility.

### Single cell RNA sequencing (scRNA-seq)

Cetuximab and untreated HNSCC cell lines were trypsinized, washed and resuspended in PBS. Cell counts and viability were made using Trypan Blue staining (ThermoFisher) in the hemacytometer. Single cell RNA labelling and library preparations were performed using the 10X Genomics Chromium^™^ Single Cell system and Chromium^™^ Single Cell 3’ Library & Gel Bead Kit v2 (10X Genomics), following manufacturer’s instructions. An input of 8,700 was used to recover a total of 5,000 cells. Sequencing was performed using the HiSeq platform (Illumina) for 2X100bp sequencing and ~50,000 reads per cell. Samples were sequenced in duplicate. Sequences were filtered and aligned using the CellRanger software (10X Genomics). Data normalization, pre-processing, dimensionality reduction (method: UMAP), cell clustering (method: louvain), differential expression analysis and visualization was performed using Monocle 3 alpha (version 2.10.1).

The scRNA-seq data are available at GEO (GSE137524)

### RNA isolation and RNA-sequencing

RNA isolation and sequencing were performed from day 0 to 5 of treatment at the Johns Hopkins Medical Institutions Deep Sequencing & Microarray Core Facility. Total RNA was isolated from at least 1000 cells collected on 1ml of QIazol reagent (Qiagen), following manufacturer’s instructions. Concentration and quality were measured at the 2100 Bioanalyzer (Agilent), with RNA Integrity Number (RIN) of 7.0 as the minimum threshold. Library preparation used the TrueSeq Stranded Total RNAseq Poly A1 Gold Kit (Illumina), according to manufacturer’s recommendations, followed by mRNA enrichment using poly(A) enrichment for ribosomal RNA removal. Sequencing was performed using the HiSeq platform (Illumina) for 2X100bp sequencing. Transcript abundance from the RNA-seq reads was inferred using Salmon ^20^. To import Salmon outputs and export into estimated count matrices we used tximport ^21^. DESeq2 was used for differential expression analysis.

All RNA-seq data are available at GEO (GSE114375).

### ATAC-sequencing

ATAC-seq library preparation was performed as previously described ^22^. Cells were collected after 5 days of treatment (100,000 cells for each group) by scrapping and were washed and lysed. Nuclei tagmentation and adapter ligation by Tn5 was performed using the Nextera DNA Sample Preparation kit (Illumina), followed by purification with MinElute PCR Purification kit (QIagen) according to manufacturers’ instructions. Transposed DNA fragments were amplified using the NEBNext Q5 HotStart HiFi PCR Master Mix with regular forward and reverse barcoded primers. Additional number of amplification cycles were determined by quantitative-PCR using the NEBNext HiFi Master Mix, SYBR Green I (Applied Biosystems) and Custom Nextera Primers. The final product was purified with MinElute PCR Purification kit (QIagen) and quality checked on 2100 Bioanalyzer (Agilent). Sequencing was performed using the HiSeq platform (Illumina) for 2X50 bp sequencing with ~50 million reads per sample.

Sequences quality were assessed using FastQC ^23^. After adapters trimming with Trim Galore! (version 0.5.0), sequences were aligned with Bowtie2 (version 2.3.2) to the human genome (hg19) ^24^. Duplicated and mitochondrial reads were removed with Picard Tools (version 2.18) ^25^, while unmapped and low quality reads were removed with Samtools (version 1.9) ^26^. MACS2 was used for peaks calling ^27^. Correlation analysis and differential bound site analysis were performed with DiffBind ^28^. The annotated differential binding sites were filtered for peaks in promoter regions.

All ATAC-seq data are available at GEO (GSE135604).

### TFAP2A RNA interference assay

Cells were transfected with a pool of four siRNA sequences (ON-TARGETplus Human TFAP2A pool, Dharmacon) to silence TFAP2A expression one day after plating. ON-TARGETplus Non-targeting Pool (NTP) and ON-TARGETplus GAPD Control Pool were used as negative and positive transfection controls, respectively. Transfection was performed in serum-free Opti-MEM (Invitrogen) and RNAiMAX Lipofectamine Reagent (Invitrogen). Eight hours after transfection, opti-MEM was replaced with complete medium and cells were incubated overnight at 37°C. Treatment with cetuximab, JQ1, the combination or vehicle was performed daily for 5 days. Transfection efficiency and level of the endogenous gene were monitored by qRT-PCR before and 72h after transfection. Cell proliferation was measured by the alamarBlue assay as described above. Each assay was performed in quintuplicate for each cell line and treatment.

### qRT-PCR analysis

Cell lines were lysed directly in the cell culture plate by adding Qiazol reagent (Qiagen) and RNA isolation followed manufacturer’s instructions. Reverse transcription of 300ng total RNA was performed with qScript Master Mix (Quanta Bioscience). Gene expression was determined using TaqMan Universal Master Mix II and TaqMan 20X Gene Expression Assays in a 7900HT equipment (Life Technologies). All assays were quantified in triplicate relative to GAPDH using the 2^-ΔΔ*Ct*^ method.

### Migration assay

The migration assays were performed in the Culture-Insert 2 Well 24 (Ibidi). In each insert well 10,000 cells (transfected and not transfected with TFAP2A siRNA) were plated and 24hs after plating treated with cetuximab, JQ1, the combination or vehicle. Once, cells were confluent the inserts were removed and gap closure was measured under a microscope at 0h, 6hs, 12hs, and 24hs. The gap area measurements were made using ImageJ 29 and closure was determined as the ratio between the initial area and the measured area at each time point. Experiments were performed at least three times.

## ACKNOWLEDGEMENTS

We thank the SKCCC Experimental and Computational Genomics Core, the JHMI Deep Sequencing & Microarray Core and the Genomic Resources Core Facility (JHU) on performing and providing advice on scRNA-seq, RNA-seq and ATAC-seq, respectively; W. Timp for advice on the ATAC-seq library preparations.

## DATA AVAILABILITY AND MATERIALS

Unless otherwise specified, all genomics analyses were performed in R. All transcriptional and epigenetic data of the cell lines from this paper have been deposited in GEO (GSE137524, GSE114375 and GSE135604).

## AUTHOR’S CONTRIBUTIONS

LTK, DAG, and EJF planned, designed and wrote the manuscript with input from all authors. LTK, FZ, JA, VY and EJF contributed to the development of methodology. LTK, MC, DAG and EJF performed analysis and interpretation of data (e.g. computational analysis). LTK, FZ, JAM VY and DAG participated in development of methodology and provided technical and material support. All authors participated in the review and/or revision of the manuscript. All authors discussed the data and contributed to the manuscript preparation. DAG and EJF instigated and supervised the project. All authors read and approved the final manuscript.

## REFERENCES

1. Sawyers, C. Targeted cancer therapy. Nature 432, 294–297 (2004).

2. Hyman, D. M., Taylor, B. S. & Baselga, J. Implementing Genome-Driven Oncology. Cell 168, 584–599 (2017).

3. Engelman, J. A. et al. MET Amplification Leads to Gefitinib Resistance in Lung Cancer by Activating ERBB3 Signaling. Science 316, 1039–1043 (2007).

4. Hata, A. N. et al. Tumor cells can follow distinct evolutionary paths to become resistant to epidermal growth factor receptor inhibition. Nature Medicine 22, 262–269 (2016).

5. Neel, D. S. & Bivona, T. G. Resistance is futile: overcoming resistance to targeted therapies in lung adenocarcinoma. npj Precision Oncology 1, (2017).

6. Shaffer, S. M. et al. Rare cell variability and drug-induced reprogramming as a mode of cancer drug resistance. Nature 546, 431–435 (2017).

7. Brand, T. M., Iida, M. & Wheeler, D. L. Molecular mechanisms of resistance to the EGFR monoclonal antibody cetuximab. Cancer Biol. Ther. 11, 777–792 (2011).

8. Vincenzi, B., Zoccoli, A., Pantano, F., Venditti, O. & Galluzzo, S. Cetuximab: from bench to bedside. Curr Cancer Drug Targets 10, 80–95 (2010).

9. Grandis, J. R. & Tweardy, D. J. Elevated levels of transforming growth factor alpha and epidermal growth factor receptor messenger RNA are early markers of carcinogenesis in head and neck cancer. Cancer Res. 53, 3579–3584 (1993).

10. Grandis, J. R. et al. Levels of TGF-α and EGFR Protein in Head and Neck Squamous Cell Carcinoma and Patient Survival. JNCI: Journal of the National Cancer Institute 90, 824–832 (1998).

11. Boeckx, C. et al. Mutation analysis of genes in the EGFR pathway in Head and Neck cancer patients: implications for anti-EGFR treatment response. BMC Res Notes 7, 337 (2014).

12. Fertig, E. J. et al. CoGAPS matrix factorization algorithm identifies transcriptional changes in AP-2alpha target genes in feedback from therapeutic inhibition of the EGFR network. Oncotarget (2016) doi:10.18632/oncotarget.12075.

13. Schmitz, S., Bindea, G., Albu, R. I., Mlecnik, B. & Machiels, J.-P. Cetuximab promotes epithelial to mesenchymal transition and cancer associated fibroblasts in patients with head and neck cancer. Oncotarget 6, 34288–34299 (2015).

14. Matys, V. et al. TRANSFAC: transcriptional regulation, from patterns to profiles. Nucleic Acids Res. 31, 374–378 (2003).

15. Matys, V. TRANSFAC(R) and its module TRANSCompel(R): transcriptional gene regulation in eukaryotes. Nucleic Acids Research 34, D108–D110 (2006). n

16. Cheng, H. et al. Decreased SMAD4 expression is associated with induction of epithelial-to-mesenchymal transition and cetuximab resistance in head and neck squamous cell carcinoma. Cancer Biol. Ther. 16, 1252–1258 (2015).

17. Singh, A. & Settleman, J. EMT, cancer stem cells and drug resistance: an emerging axis of evil in the war on cancer. Oncogene 29, 4741–4751 (2010).

18. Fuchs, B. C. et al. Epithelial-to-mesenchymal transition and integrin-linked kinase mediate sensitivity to epidermal growth factor receptor inhibition in human hepatoma cells. Cancer Res. 68, 2391–2399 (2008).

19. Hafner, M., Niepel, M., Chung, M. & Sorger, P. K. Growth rate inhibition metrics correct for confounders in measuring sensitivity to cancer drugs. Nature Methods 13, 521–527 (2016).

20. Patro, R., Duggal, G., Love, M. I., Irizarry, R. A. & Kingsford, C. Salmon provides fast and bias-aware quantification of transcript expression. Nat Methods 14, 417–419 (2017).

21. Soneson, C., Love, M. I. & Robinson, M. D. Differential analyses for RNA-seq: transcript-level estimates improve gene-level inferences. F1000Res 4, 1521 (2015).

22. Buenrostro, J. D., Giresi, P. G., Zaba, L. C., Chang, H. Y. & Greenleaf, W. J. Transposition of native chromatin for fast and sensitive epigenomic profiling of open chromatin, DNA-binding proteins and nucleosome position. Nat. Methods 10, 1213–1218 (2013).

23. Andrews, S. FastQC: a quality control tool for high throughput sequence data. (2010).

24. Langmead, B. & Salzberg, S. L. Fast gapped-read alignment with Bowtie 2. Nat Methods 9, 357–359 (2012).

25. Wysoker, A., Tibbetts, K. & Fennell, T. Picard Tools Version 2.18. http://broadinstitute.github.io/picard/.

26. Li, H. et al. The Sequence Alignment/Map format and SAMtools. Bioinformatics 25, 2078–2079 (2009).

27. Zhang, Y., Liu, T., Meyer, C. A. & Eeckhoute, J. Model-based analysis of ChIP-Seq (MACS) Genome Biol. 2008; 9 (9): R137. (2008).

28. Ross-Innes, C. S. et al. Differential oestrogen receptor binding is associated with clinical outcome in breast cancer. Nature (2012) doi:10.1038/nature10730.

29. Schneider, C. A., Rasband, W. S. & Eliceiri, K. W. NIH Image to ImageJ: 25 years of image analysis. Nat. Methods 9, 671–675 (2012).

30. Maseki, S. et al. Efficacy of gemcitabine and cetuximab combination treatment in head and neck squamous cell carcinoma. Mol Clin Oncol 1, 918–924 (2013).

31. Leonard, B. et al. BET Inhibition Overcomes Receptor Tyrosine Kinase-Mediated Cetuximab Resistance in HNSCC. Cancer Res. 78, 4331–4343 (2018).

32. Schulte, J. H. et al. Transcription factor AP2alpha (TFAP2a) regulates differentiation and proliferation of neuroblastoma cells. Cancer Letters 271, 56–63 (2008).

33. Ding, X. et al. Transcription factor AP-2α regulates acute myeloid leukemia cell proliferation by influencing Hoxa gene expression. Int. J. Biochem. Cell Biol. 45, 1647–1656 (2013).

34. Carrière, C., Mirocha, S., Deharvengt, S., Gunn, J. R. & Korc, M. Aberrant expressions of AP-2α splice variants in pancreatic cancer. Pancreas 40, 695–700 (2011).

35. Chambers, S. M. et al. Highly efficient neural conversion of human ES and iPS cells by dual inhibition of SMAD signaling. Nat Biotechnol 27, 275–280 (2009).

36. Basu, D. et al. EGFR inhibition promotes an aggressive invasion pattern mediated by mesenchymal-like tumor cells within squamous cell carcinomas. Mol. Cancer Ther. 12, 2176–2186 (2013).

37. Byers, L. A. et al. An epithelial-mesenchymal transition gene signature predicts resistance to EGFR and PI3K inhibitors and identifies Axl as a therapeutic target for overcoming EGFR inhibitor resistance. Clin. Cancer Res. 19, 279–290 (2013).

38. Brand, T. M. et al. AXL mediates resistance to cetuximab therapy. Cancer Res. 74, 5152–5164 (2014).

39. Zhang, Z. et al. Activation of the AXL kinase causes resistance to EGFR-targeted therapy in lung cancer. Nat. Genet. 44, 852–860 (2012).

40. Filippakopoulos, P. et al. Selective inhibition of BET bromodomains. Nature 468, 1067–1073 (2010).

41. Decker, T.-M. et al. Transcriptome analysis of dominant-negative Brd4 mutants identifies Brd4-specific target genes of small molecule inhibitor JQ1. Sci Rep 7, 1684 (2017).

42. Stratikopoulos, E. E. et al. Kinase and BET Inhibitors Together Clamp Inhibition of PI3K Signaling and Overcome Resistance to Therapy. Cancer Cell 27, 837–851 (2015).

43. Alqahtani, A. et al. Bromodomain and extra-terminal motif inhibitors: a review of preclinical and clinical advances in cancer therapy. Future Sci OA 5, FSO372 (2019).

44. Andrieu, G., Belkina, A. C. & Denis, G. V. Clinical trials for BET inhibitors run ahead of the science. Drug Discov Today Technol 19, 45–50 (2016).

