## Supplementary figures for "Integrated single cell and bulk gene expression and ATAC-seq reveals heterogeneity and early changes in pathways associated with resistance to cetuximab in HNSCC sensitive cell lines"

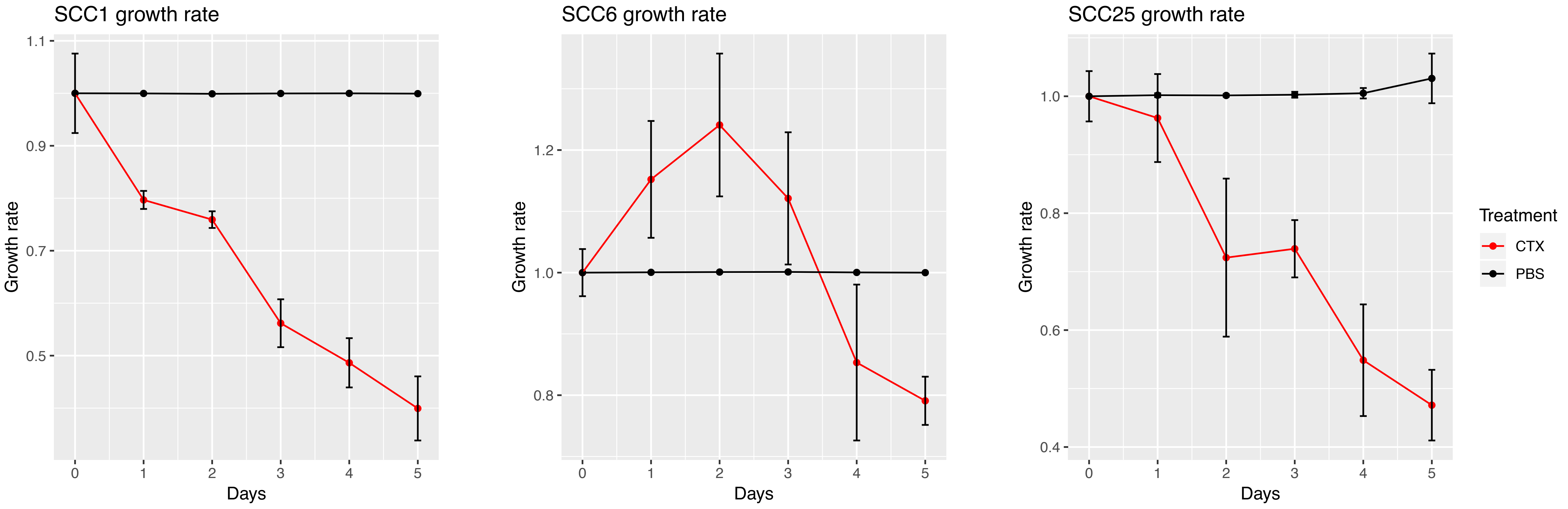


**Supplementary Figure 1 -** Proliferation assay to verify SCC1, SCC6 and SCC25 sensitivity to cetuximab. HNSCC cells were seeded in 6-well plates and treated with cetuximab or PBS daily for five days. Proliferation was measured every 24 hours using the alamar blue assay. The fluorescence measured was used to determine the growth rate ($GR = 2k^{(c,t)/k(0)} - 1$).

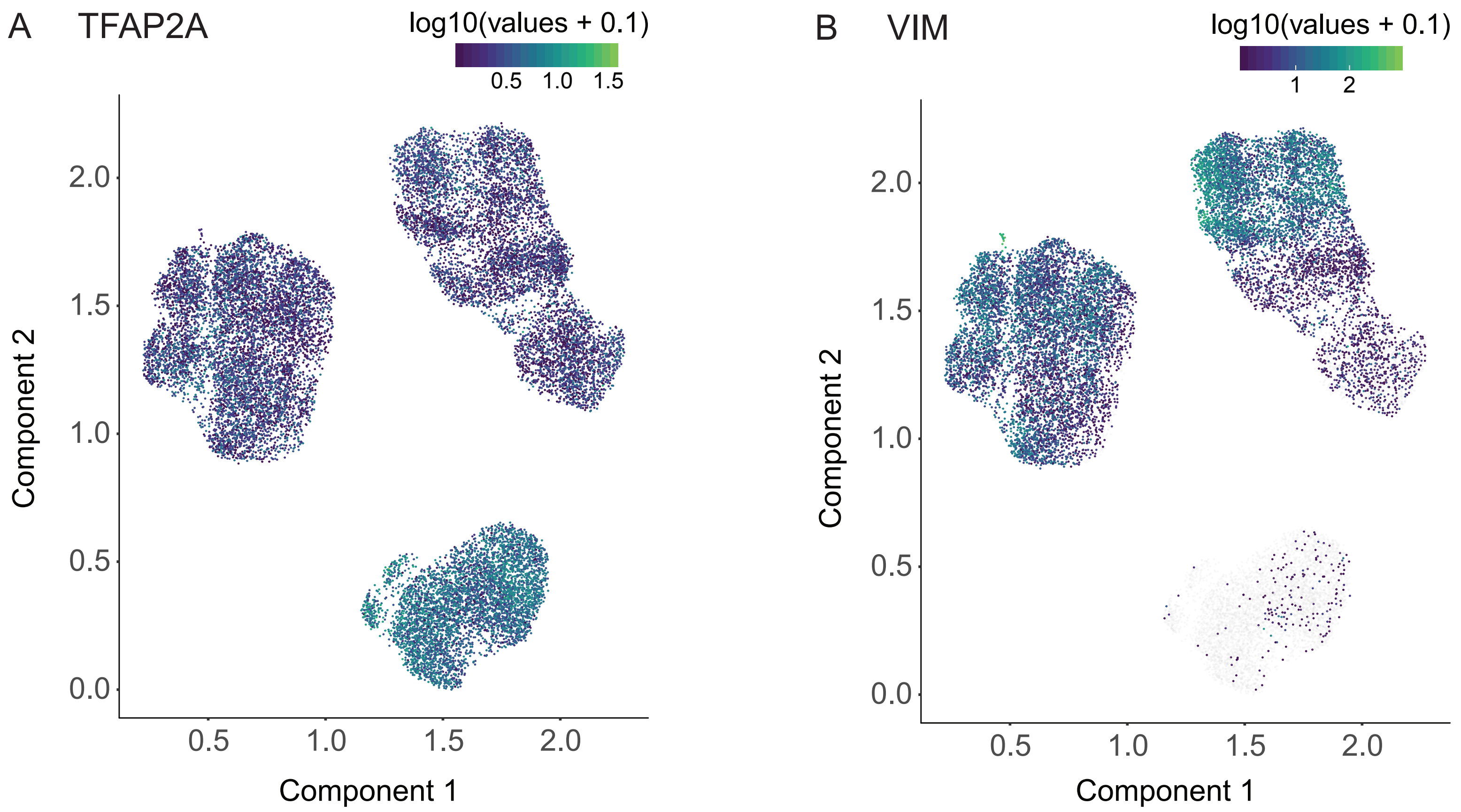


**Supplementary Figure 2 -** Single cell expression of TFAP2A and VIM in SCC1, SCC6 and SCC25.

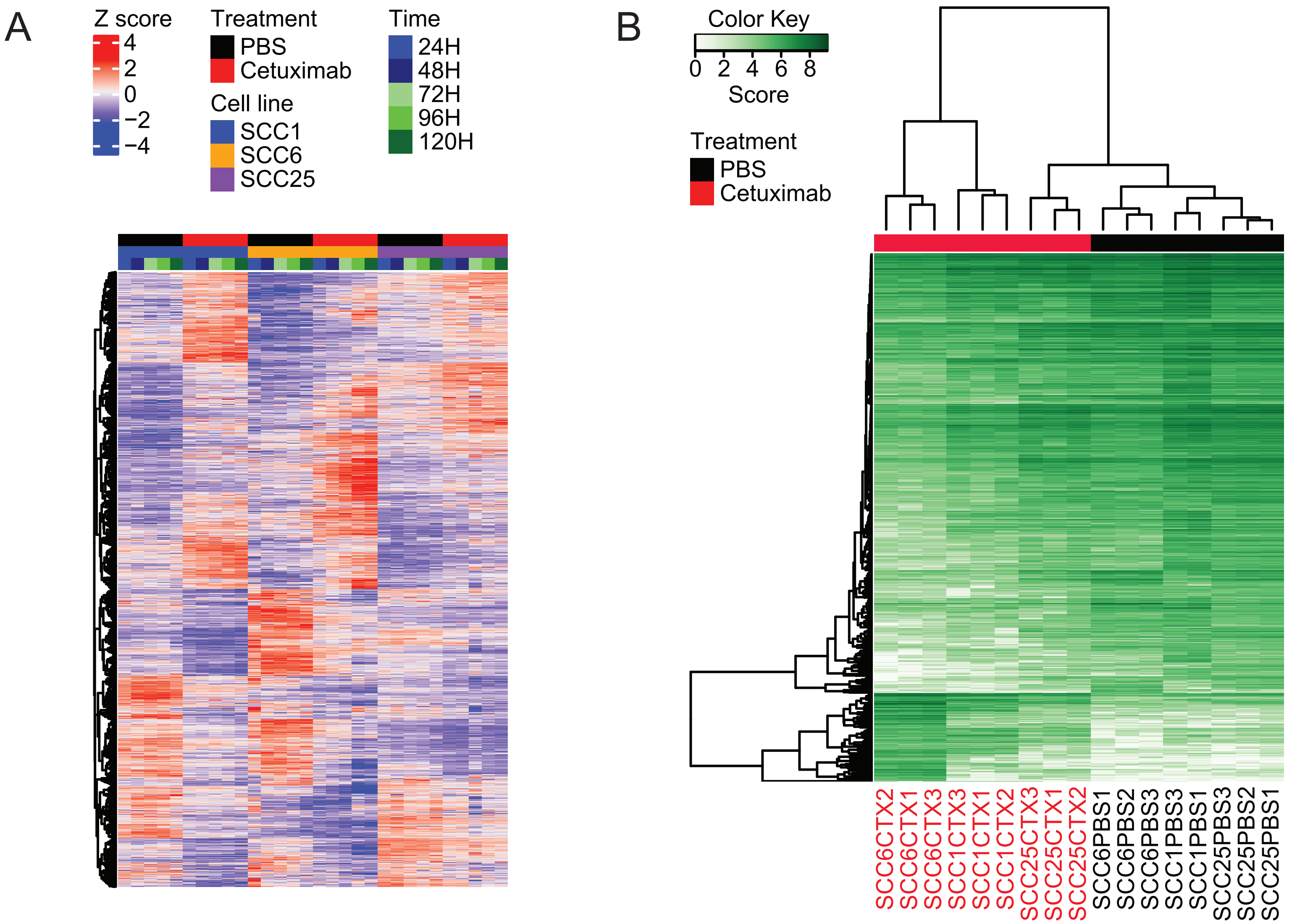


**Supplementary Figure 3 -** Differential expression analysis (A) and differential binding analysis (B) for all SCC1, SCC6 and SCC25 cell lines. RNA-seq analysis for differentially expressed genes was performed to identify the common genes that change overtime in all three cell lines. Differential binding analysis on the ATAC-seq peaks was also performed in all three cell lines at the same time to identify the common bound sites with accessibility changes in response to cetuximab therapy.
